# *Rab11* is essential for *lgl* mediated JNK–Dpp signaling in dorsal closure and epithelial morphogenesis in *Drosophila*

**DOI:** 10.1101/713115

**Authors:** Nabarun Nandy, Jagat Kumar Roy

## Abstract

Dorsal closure in *Drosophila* provides a robust genetic platform providing deep insights into the basic cellular mechanisms that govern epithelial wound healing and morphogenesis. As dorsal closure proceeds, the adjacent epithelia advance contra-laterally involving coordinated cell shape changes in order to successfully accomplish the process. The JNK-Dpp signaling in these cells plays an instrumental role in guiding their fate as gastrulation completes. A huge number of genes have been reported to be involved in the regulation of this core signaling pathway, yet the mechanisms by which they do so is hitherto unclear, which forms the objective of our present study. Here we show that *lgl*, which is a potent tumour suppressor gene, conserved across the phyla till humans, regulates the JNK–Dpp pathway in the dorsal closure and epithelial morphogenesis process where in ectopic knockdown of this gene results in the failure of dorsal closure. Interestingly, we also find *Rab11* to be interacting with *lgl* as they together regulate the core JNK-Dpp signaling pathway during dorsal closure and also during pupal thorax closure process. Using the robust *Gal4-UAS* system of targeted gene expression, we show here that *Rab11* and *lgl* synergize to successfully execute the dorsal closure and the similar thorax closure process, ensuring proper spatio-temporal JNK-Dpp signaling.

## Introduction

The spectrum of cellular mechanisms employed by tumour suppressor mutations in order to help tumours grow and disseminate is diverse and enigmatic. A vast proportion of these tumour suppressor genes comprise of the ones coding for cell polarity (Bilder et al, 2000; Humbert et al,2003; Royer and Lu, 2011), which also happen to be developmentally active genes (Klezovitch et al, 2004) as they determine cellular differentiation at the time of tissue remodeling and morphogenesis (Tanentzapf and Tepass, 2003). Cell polarity determining genes, like other tumour suppressor genes, are developmentally active and exert their phenotype in the homozygous recessive state. Cellular polarity represents its differentiated state, especially in an epithelial tissue where the constituent cells show a robust apico-basal polarity and planar polarity. These polarities are determined by proteins or protein complexes which occupy distinct domains, either on the cell membrane or in the cytoplasm and their dynamics and cross talk with intracellular signaling bring about different cellular changes necessary for shaping tissues at the time of development. One such polarity determining gene is *lgl* which not only is a developmentally active gene but is also a potent tumour suppressor, capable of organizing intracellular cyto-architecture on one hand and on the other hand is responsible for regulating critically important cell signaling pathways like JNK (Zhu et al, 2010), Hippo (Sun and Irvine, 2011; Enomoto and Igaki, 2011) and Notch (Parsons et al, 2014, Langevin et al, 2005; Portela et al, 2015). These signaling pathways have been studied in the context of the classically acclaimed function of *lgl* which is tumorigenesis (Humbert et al, 2008, Hariharan and Bilder, 2006). Other than this function *lgl* has also been reported to be an essential regulator of neurogenesis (Peng et al, 2000) which if perturbed, can give rise to Brain tumours.

The early reports of *lgl’s* involvement in the process of embryonic epithelial morphogenesis arrives from the reports of Manfruelli et al, 1996, where temperature sensitive hypomorph alleles of *lgl* mutants show dorsal closure and head involution defects when reared at elevated temperatures. Similar observations were made by Tanentzapf and Tepass, 2002; Hutterer et al, 2004, where the latter group has revealed the interactions between the CDC-42, PAR6, APKC and *lgl* in the establishment of epithelial polarity in the developing embryos. Here it has been proven that PAR-6 protein’s apical localization is CDC-42 dependent whereas on the other hand its expulsion from the baso-lateral cell cortex is *lgl* dependent. *lgl* activity is further facilitated by the activity of APKC which phosphorylates *lgl* in the apical cytoplasm and restricts its activity in the basal domain of the differentiated epithelial cells. Bilder et al, 2000, have shown the interaction of *lgl* with the junctional proteins, Scrib and Dlg, which form a strongly interacting Trio, indispensable for the establishment of epithelial cell polarity. Scrib and Dlg molecules remain physically associated with each other and they ascertain Lgl’s presence and activity in the baso-lateral domain. The mutants of all three genes thus have similar consequences as all three of them behave as potentially strong tumour suppressors.

The activity of *lgl* has been shown to be intimately associated with the proper execution of the JNK pathway as has been reported by Zhu et al, 2010. The JNK signaling pathway is a core signaling pathway in the process of dorsal closure at the time of gastrulation in *Drosophila* embryos (Noselli, 1998; Noselli and Agnes, 1999; Ramet et al 2002; Stronach and Perimon, 2002). According to the reports of Noselli and Agnes, 1999, a strong JNK expression in the dorsal most lateral epithelial cells of the dorso-lateral epidermis (DLE) leads to the downstream expression and secretion of the Dpp morphogens under whose influence the dorsolateral epithelial cells undergo coordinated elongations in order to bring about the contralateral movement of the lateral epithelia (LE) which ultimately zippers the dorsal opening. It has been observed by Arquier et al, 2001 that *lgl* plays an instrumental role in the process of release of Dpp morphogen via exocytosis and acts upstream of its receptor *Thickveins.* This observation finds support from the studies of Zhang et al, 2005 where they have reported in a Yeast model that Lgl mediates polarized exocytosis by exerting its influence on the exocyst complex which tethers intracellular vesicles as they subsequently get docked to their cognate SNAREs on the target membranes. In this light, *lgl* also seems to be regulating intracellular vesicle transport which forms an essential mode of directed trafficking amidst the vast network of endomembrane systems present inside eukaryotic cells. An essential component of this endomembrane system is the recycling endosomes which participate in membrane recycling and exocytosis and is essentially marked with Rab11, a small conserved Ras like GTPase, which remains associated with vesicles emanating and fusing between recycling endosomes, trans Golgi apparatus and plasma membrane.

The present study focuses on the interaction of the tumour suppressor *lgl* with *Rab11* owing to its property to promote polarized trafficking inside differentiated cells. Rab11 has been proven beyond doubt to be interacting with essential components of the exocyst complex like Sec15 (Wu et al, 2005; Langevin et al, 2005; Guichard et al, 2015; Bhuin and Roy, 2019) and Nuf (Cao et al, 2008). Here we find that a targeted and conditional down-regulation of *lgl* using the robust *Gal4-UAS* (Brand and Perrimon,1993) system of targeted gene expression in *Drosophila*, in the dorsolateral epithelium of the gastrulating fly embryos, result in an up-regulation Rab11 expression, which affects normal JNK-Dpp signaling in the DLE of the embryos. A down-regulation of Rab11 in the *lgl* down-regulated genetic background results in a significant rescue of the *lgl* down-regulated phenotype and a consequent restoration of the regular JNK and Dpp signaling pattern, which suggests a strong genetic interaction between the two.

## Materials and Methods

### Fly stocks and rearing conditions

All fly stocks have been reared on standard food preparation containing maize powder, agar, yeast and sugar with methyl-p-hydroxy benzoate as anti-fungal and also propionic acid as anti-bacterial agents at a temperature of 23±1°C. The stock *pnr*^*MD237*^/*TM3,Ser* was obtained from Bloomington *Drosophila* Stock Center (BDSC 3039, Thomas et al, 2009) and expresses Gal4 as reported by Calleja et al, 1996, which was further introgressed with *TM3, ActGFP, Ser*^1^/*TM6B* in order to generate *pnr*^*MD237*^/*TM3, ActGFP, Ser*^1^stock. *TRE-JNK* (Chatterjee and Bohmann, 2012) was introgressed with *Sp/CyO; pnr*^*MD237*^/ *TM3, ActGFP, Ser*^1^ and *Sp/CyO; pnr*^*MD237*^/*TM6B* to obtain *TRE-JNK/CyO*; *pnr*^*MD237*^/*TM3*, *ActGFP, Ser*^1^ and *TRE-JNK/CyO*; *pnr*^*MD237*^/*TM6B* stocks, respectively. *UAS-Rab11*^*RNAi*^; + (Satoh et al, 2005), was introgressed with *+; UAS-lgl*^*RNAi*^ (BDSC 35773, Perkins et al 2015) to obtain *UAS-Rab11*^*RNAi*^; *UAS-lgl*^*RNAi*^ stocks. *UAS-YFP-Rab11*^*Q70L*^ stock (Zhang et al, 2007) was similarly introgressed with *UAS-lgl*^*RNAi*^ in order to obtain *UAS-YFP-Rab11*^*Q70L*^/*CyO, ActGFP;UAS-lgl*^*RNAi*^ stock. *Dpp-lacZ/CyO* (BDSC 68153) flies were double balanced to obtain *Dpp-LacZ/CyO-Act-GFP; dco2/ TM3, ActGFP, Ser*^1^ stock which was further introgressed with *Sp/CyO-Act-GFP; pnr*^*MD237*^/ *TM3, ActGFP, Ser*^1^ to obtain *Dpp-LacZ/CyO, ActGFP; pnr*^*MD237*^/*TM3, ActGFP, Ser*^1^ stock. This stock was used as a driver in our experiments where only non-GFP embryos obtained from the crosses set with the *UAS* alleles of *Rab11* and *lgl* were proceeded for β-galactosidase staining.

### Embryo Collection and Fixation

All flies were made to lay eggs on standard agar plates supplemented with sugar and propanoic acid and eggs were collected as per the protocol suggested by Narasimha and Brown, 2006, with slight modifications. For whole mount preparations and immunostaining of embryos, different alleles and transgenes were balanced with GFP tagged balancers and only non GFP or driven embryos were selected for experimental purpose. Eversed clypaeolabrum was treated as a marker of stage 13 and retracted clypaeolabrum was treated as marker of stage 14 (Sasikumar and Roy, 2009). Embryo staging was done according to Hartenstein’s Atlas of *Drosophila* Development, 1993.

### Genetics

In order to observe the tissue specific effects of *Rab11*, the *Gal4-UAS* system as described by Brand and Perrimon, 1993, was used to drive its alleles in stage13 and 14 fly embryos. *pnr^MD237^/ TM3, ActGFP, Ser* stock was used as the Gal4 driver and *UAS* constructs of different alleles of *Rab11* were used to observe tissue specific effects of these genes. Males of *Gal4* stock and Virgin Females of *UAS* constructs were used to set up crosses in order to obtain embryos of the desired genotype. The desired genotypes were screened on the basis of GFP expression of the balancer chromosomes (Supplementary figure S1). These embryos were further staged as suggested by Green et al, 1993. The embryos of identical stages were further used for analysis of cell-biological and molecular parameters.

### Embryonic cuticle preparations

*Drosophila* embryonic cuticles were prepared as described by Anderson, 1985; Ostrowski, 2002 with slight modifications. The cuticles were fixed in glycerol:acetic acid (1:4) solution for 60 min at 37°C, mounted in Hoyer’s mounting medium and then baked overnight (∼12-14h) at 65°C. These cuticles were subjected to dark field microscopy in Nikon eclipse E800 microscope, and the images obtained were further processed using the aid of the Adobe Photoshop CS6 software. The embryonic cuticle of *Drosophila* has been extensively used to study the morphogenesis of the underlying epidermis. Any defect of the underlying epidermis thus becomes fairly evident in the secreted cuticle which can be observed by the above mentioned technique.

### Immunostaining, imaging and confocal microscopy

*Drosophila* embryos were fixed and imaged as described by Narasimha and Brown, 2006. The dechorionated and devitellized embryos were fixed in 4% para-formaldehyde solution and stored in absolute methanol. For immunostaining, these embryos were rehydrated using methanol gradients of 70%, 50%, 30% and 10% in 0.1% PBT solution. The embryos were blocked for 2h at RT in blocking solution as described by Banerjee and Roy, 2017. Rabbit anti-sera against *Drosophila* Rab11 (Alone et al, 2005) was used at a dilution of 1:100 for immunostaining and 1:1000 for western, DSHB anti-FasIII (7G10) antibody was used and secondary antibodies were used as described by Sasikumar and Roy, 2009; Bhuin and Roy 2012; Ray and Lakhotia, 2017, and were imaged using single photon confocal microscope using Zen software, 2012. The images obtained were analyzed using Zeiss LSM 510 Meta-software.

Dark field, fluorescence and phase contrast images of the embryos were taken under the Nikon eclipse E800 microscope under the same gain and exposure values. 22-24h developed embryos were dechorionated and mounted in halocarbon oil in order to image them live in phase contrast as well as fluorescence microscope.

### Embryonic, pupal and larval lethality assays were performed according to standard procedures

The *Gal4-UAS* system of targeted gene expression was used in order to see the effects of alleles of *Rab11* on the embryonic lethality, where in every experiment males of the *Gal4* and virgin females of the *UAS* constructs were introgressed and embryos were collected from the F1 generation. These embryos were incubated for 24 to 48 h at 23°C on standard agar plates and the total number of dead embryos (detected by yellowing colour of the eggs) were counted against total number of fertilized eggs. These fertilized eggs include the dead as well as the hatched embryos, and percentage death was calculated as:

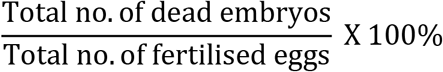

The % lethality for each cross was calculated in triplicates and the mean lethality so obtained was tabulated and graphically represented using MS-Excel-2013 spreadsheet software. The lethality caused due to balancers has been subtracted from those introgressions which involve balancer chromosomes in the *Gal4* and the *UAS* stocks.

The final percentages have been calculated as explained in the following instances:

In a *pnr-Gal4/TM3, Ser* introgressed with *UAS-Rab11*^*N124I*^/*CyO* experiment, *pnr-Gal4/TM3, Ser* introgressed with *+/CyO* has been taken as a control. The lethality observed in the latter case was subtracted from the lethality observed in the experimental case. The final lethality % value was multiplied by 4 as only 1/4^th^ of the F1 obtained from the experimental cross would have *UAS-Rab11*^*N124I*^ driven by *pnr-Gal4* according to Mendelian ratios. Similarly if *UAS-Rab11*^*RNAi*^ were to be driven by the same *Gal4*, the control cross would be *pnr-Gal4* driven +/+. The lethality or eclosion percentage obtained in the latter would be subtracted from the experimental cross and the resulting value would be multiplied by 2 as only 50% of the progeny derived from the experimental cross would have *UAS-Rab11*^*RNAi*^ driven by *pnr-Gal4*.

### β-galactosidase (LacZ) reporter assay

In order to observe the Dpp expression pattern in a mutation deficient background, *dpp-LacZ/CyO* (Bloomington stock: 68153) flies were brought in a wild type background by introgression with *+/+* flies such that the F1 progeny had the *dpp-LacZ/+* genotype, which were then made to lay eggs on standard sugar-agar plates from which eggs were collected, dechorionated and devitellized according to standard protocol described by Sasikumar and Roy, 2009. These embryos were rinsed thoroughly in 1X PBS and fixed in 4% PFA for 10 min. After thorough rinsing with 1X PBS these were then suspended in LacZ staining solution with 8% X-Gal in DMSO pre-incubated at 37°C for 1h or till colour developed. As the embryos developed blue colour, they were washed in 1X PBS solution and mounted in 70% Glycerol in Bridge slides. The same protocol was followed in the experimental conditions also. The embryos were imaged under a Nikon Digital Sight DS-Fi2 camera installed on a Nikon SMZ800N stereo-binocular microscope.

### Semi-Quantitative RT-PCR

12-13h synchronized, 200 embryos of stage 14-early stage 15 were collected, washed in DEPC treated 1X-PBS and then proceeded for total RNA isolation by the Trizol method as prescribed by the manufacturer’ protocol (Sigma Aldrich, India). RNA pellets so obtained were dissolved in nuclease free MQ water and quantified spectrophotometrically. 1 µg of RNA from these samples were incubated with 2U of RNase free DNaseI (MBI, Fermentas, USA) for 30 min at 37°C for removal of any remaining DNA. First strand cDNA synthesis was carried out by 1 µg of total RNA as described by Ray et al, 2019. The prepared cDNA were subjected to PCR amplification for a 25µl of reaction mix containing 50 ng of cDNA, 25pM each of the forward and reverse primers, 200µM of each dNTP (Sigma Aldrich USA) and 1.5U of Taq DNA Polymerase (Geneaid, Bangalore), which were carried out under the following conditions: Denaturation for 3min at 94°C followed by annealing for 30 sec at 60°C with an extension for 30 sec at 72°C for 30 cycles and with a 7 min final extension at 72°C in the last cycle. The products were run on a 2% Agarose gel with a 50 bp ladder. The following were the primer sequences for *lgl* and *G3PDH*:

*lgl* (forward and reverse):

ATAGAGATGTCGCTGAAGTTCTTGT

GAGTGAAGATATGGCGCTTTGATAG

*G3PDH* (forward and reverse):

CCACTGCCGAGGAGGTCAACTA

GCTCAGGGTGATTGCGTATGCA

Gel images were analyzed in UV Transilluminator gel documentation and analysis system (Syngene). For all RT–PCR analysis, band intensities were measured by two methods: Alpha imager software and Histogram tool of Adobe photoshop CS6 software.Each experiment was carried out four times from which mean values were calculated taking into account the slightest variations.

## Results

### A targeted down-regulation of *lgl* in the dorso-lateral epidermis resulted in the dorsal open phenotype similar to JNK pathway mutants

The embryonic cuticle of *Drosophila* is an index of proper epithelial morphogenesis process involving a large amount of tissue level and cell biological changes which at the time of development is regulated by a large number of genetic and environmental factors. Any perturbation in these factors leads to an improper morphogenesis process which is detectable in the secreted cuticle, thereby making it an excellent indicator of the successful execution of morphogenesis process. Thus 22-24 h synchronized *pnr-Gal4* (*pnr*^*MD237*^) driven *UAS-lgl*^*RNAi*^ embryos were proceeded for cuticle preparations according to standard protocols, using *pnr*^*MD237*^/+ and *+/+* embryos as controls. The cuticles were imaged by Dark Field Microscopy at 20X magnification under Nikon Eclipse E800 microscope and the final images were processed using Adobe Photoshop CS6 software. It was observed that *pnr-Gal4* driven *UAS-lgl*^*RNAi*^ embryos show a characteristic puckering of the cuticles along with a characteristic dorsal opening where 23 out of a total 66 cuticles observed, showed this phenotype when embryos were collected from the F1 of a cross between *pnr*^*MD237*^/*TM3,Act-GFP* males and *UAS-lgl*^*RNAi*^ females. These embryos also reveal a characteristic dorsal opening, much like the puckered loss of function mutants as compared to the undriven embryos which show no such features.

### Leading edge cells and cells of the dorso-lateral epidermis show significant morphological defects in *Rab11*, *basket* and *lgl* conditional mutants

Conditionally driven *UAS-bsk*^*DN*^ and *UAS-lgl*^*RNAi*^ mutants showed strong resemblances in their cuticular phenotypes which could arise due to similar cell morphological defects. Thus, in order to observe the cellular defects in all of the above genetic backgrounds, stage 14 (embryos with retracted Clypaeolabrum) driven embryos were at first screened on the basis of GFP and non GFP (Note: *pnr-Gal4* is balanced with *TM3, Ser, Act-GFP*) (Supplementary data S1) and the non-GFP embryos were proceeded for immune-staining with anti-FasIII antibody and were then counterstained with DAPI. FasIII was detected with anti-mouse AF-488 secondary and imaged under the confocal microscope. Corresponding projections and magnified sections have been shown in the alongside figure. The morphology and the shape of LE and DLE cells show a patterned elongated structure in the wild type conditions. However, the mutants lack this characteristic feature, instead they assume a somewhat hexagonal structure which signifies that the elongated morphology of the DLE cells is under the strong influence of the region specific bsk/JNK signaling, which, if perturbed results in a failure of the necessary morphological changes. Here we also show that a targeted down-regulation of *lgl* by driving *UAS-lgl*^*RNAi*^ by *pnr-Gal4* shows a similar phenotype which could be in a way regulating proper JNK-Dpp signaling in that region. The targeted down-regulation of *lgl* in *pnr-Gal4* driven *UAS-lgl*^*RNAi*^ individuals thus show a characteristic dorsal closure defect as seen in JNK pathway mutants like *basket, hep, slpr or puc*.

### *Rab11* interacts with *lgl* in the process of epithelial morphogenesis

The *lgl* gene in *Drosophila* codes for a PDZ domain containing cytoskeletal protein which is a major baso-lateral polarity determinant in differentiated epithelial cells. Apart from its strong roles in tumorigenesis, which seems to be conserved across the phyla, it is also a potentially strong developmentally active gene which has been studied and shown to regulate the process of epithelial morphogenesis (Manfruelli et al, 1996). In our experiments, we find that *lgl* shows a strong genetic interaction with Rab11. We observed this interaction as a rescue of the embryonic lethality observed in *pnr*^*MD237*^ driven *UAS-lgl*^*RNAi*^; *UAS-Rab11*^*RNAi*^ individuals as compared to the individual embryonic lethality values obtained from the F1 of *pnr-Gal4* driven *UAS-lgl*^*RNAi*^ and *pnr-Gal4* driven *UAS-Rab11*^*RNAi*^. Similarly an upsurge of embryonic lethality was observed when *UAS-Rab11CA (UAS-YFP-Rab11*^*Q70L*^) introgressed with *UAS-lgl*^*RNAi*^ was driven by *pnr-Gal4* suggesting that perturbation of *lgl* in the epithelial cells essentially affects Rab11 titers thereby suggesting its regulatory effect over intracellular Rab11 levels.

As we show in Fig 3, the dorso-ventral polarity of cells of the LE is evident from their regular rhomboid geometry, with the DLE cells showing a significant dorso-ventral elongation. This suggests their importance as they further induce cell shape changes in the dorsolateral epithelium necessary for the successful execution of the dorsal closure process. Stage 14 embryos, were stained for Zona Occludens (septate junction) marker, FasIII (green), and counterstained with DAPI. It was observed that the cells of the LE in *pnr-Gal4* driven *UAS-lgl*^*RNAi*^ individuals and *pnr-Gal4* driven *UAS-Rab11*^*RNAi*^ individuals show a considerable loss of this geometry as well as polarity, where the cells show all sorts of shapes, like hexagon, pentagon or even circles. This is a clear indicative of the fact that these cells undergo a considerable degree of polarity loss when *Rab11* or *lgl* is individually knocked down, however in the latter case, the extent of this polarity loss is more pronounced as compared to the *Rab11* knockdown individual. However, when both the genes are knocked down simultaneously, a rescue was obtained where cells more or less resembled the wild type morphology.

It was also observed that from a total of 389 *pnr-Gal4* driven *UAS-lgl*^*RNAi*^ individuals, 45 individuals or 11.56% were found dead. After subtraction of a balancer lethality of 1.39% from this observed value, net lethality turned out to be 10.18%. As this result was obtained in 50% of driven progeny, therefore out of an expected 100% driven progeny, lethality value turns out to be 20.36%. Similarly from a total of 402 *pnr-Gal4* driven *UAS-Rab11*^*RNAi*^ individuals, 37 individuals or 9.2% were found dead. After subtraction of balancer lethality of 1.39%, net lethality turned out to be 7.81%. As this result was obtained in 50% of driven progeny therefore out of an expected 100% driven progeny, lethality value was calculated as15.62%. Again, from a total of 491 *pnr-Gal4* driven *UAS-Rab11*^*RNAi*^; *UAS-lgl*^*RNAi*^ individuals, 11 individuals or 2.24% were found dead. After the subtraction of a balancer lethality of 1.39% net lethality turned out to be 0.85%. As this result was obtained in 50% of driven progeny therefore out of an expected 100% driven progeny, lethality value turns out to be 1.7%. This suggests a strong rescue of *lgl* knockdown phenotype by a simultaneous *Rab11* knockdown in the same tissue.

A complete reversal of the phenotype was seen when *UAS-Rab11*^*CA*^ allele was simultaneously driven in an *lgl*^*RNAi*^ background by *pnr-Gal4.* It was observed that from a total of 218 *pnr-Gal4* driven *UAS-Rab11*^*CA*^ individuals, 6 individuals or 2.75% were found dead. After subtraction of balancer lethality of 2.39%, net lethality turned out to be 0.36%. As this result was obtained in 25% of driven progeny therefore, out of an expected 100% driven progeny, lethality value turns out to be 1.44%. On a similar note, it was observed that from a total of 418 *pnr-Gal4* driven *UAS-Rab11*^*CA*^; *UAS-lgl*^*RNAi*^ individuals, 293 individuals or 70.09% were found dead. After subtraction of balancer lethality of 56.39%, net lethality turned out to be 13.70%. As this result was obtained in 25% of driven progeny therefore out of an expected 100% driven progeny, lethality value turns out to be 54.82%. This suggests a strong upsurge of the lethal phenotype when *Rab11* is over-expressed in an *UAS-lgl*^*RNAi*^ background. Supplementary Table 1 further elaborates our observations as it provides greater details of the cross schemes set up to study the consequence of *Rab11* and *lgl* perturbation in embryonic survivability.

### *lgl* knockdown in the dorsolateral epidermis shows a significant upsurge of *Rab11* expression levels

As shown in our previous experiments an *UAS-lgl*^*RNAi*^ or a *UAS-Rab11*^*RNAi*^ when driven by pnr-Gal4, results in dorsal closure defects much like JNK pathway mutants, provided these constructs are driven individually. However, a simultaneous down-regulation of both *UAS-lgl*^*RNAi*^ as well as *Rab11*^*RNAi*^ in the same embryos results in the rescue of the dorsal open condition which consequently manifests in the form of embryonic survivability. In order to observe the expression of Rab11 in a *pnr-Gal4* driven *UAS-lgl*^*RNAi*^ background, stage 13-14 embryos (detected by their eversed clypeolabrum) were immunostained for Rab11 and FasIII antigens and detected with anti-Rabbit AF-488 and anti-mouse AF-546 secondary antibodies. It was observed that a strong up-regulation of *Rab11* follows the knockdown of *lgl* by *pnr-Gal4* driven *UAS-lgl*^*RNAi*^ in the DLE. On driving *UAS-Rab11*^*RNAi*^; *UAS-lgl*^*RNAi*^ by *pnr-Gal4*, results in a rescue. This suggests that a Rab11 over-expression in an *lgl* knockdown background could be responsible for the *lgl* mediated phenotypes, at least as seen in the case of embryonic dorsal closure process (Fig. 4).

### *Rab11* and *lgl* show a genetic interaction in the process of thorax closure during pupal development and adult thorax formation

The events of dorsal closure and thorax closure employ overlapping cellular mechanisms such as coordinated cell shape changes in order to bring about their successful execution. These processes trigger some core signaling cascades like the JNK-Dpp pathway, whose upstream regulation is again governed by the proper expression pattern of several genes regulating different aspects of cell biology. To confirm, whether the two genes i.e., *lgl* and *Rab11* interact in a similar manner at the time of thorax closure process also, which involves the contralateral elongation and fusion of wing imaginal disc nota again under the influence of JNK-Dpp signaling.

Observation of dorsal epithelial morphologies of larva, pupa, and adult *Drosophila* were observed which suggested that the interaction of *Rab11* and *lgl* is essential for the morphogenesis of the dorsal epithelium wherein embryos which are rescued in a simultaneous *lgl* and *Rab11* knockdown condition as shown in the previous experiments, emerge as larvae with and without dorsal epithelial lesions (Fig. 5a). Out of a total of 650 non-tubby third instar larvae obtained from the cross *pnr-Gal4/TM6B* X *UAS-Rab11*^*RNAi*^; *UAS-lgl*^*RNAi*^, 566 larvae or a mean of 87.76% of the rescued third instar larvae show dorsal lesions or a partial rescue and they do not emerge as flies exhibiting drastic thorax closure defects (Fig. 5b), whereas the ones with complete rescue ∼12.23% do not show any dorsal closure defects and emerge as healthy flies (Fig. 5d).

Fly eclosion data was also calculated for *pnr-Gal4* driven *UAS-lgl*^*RNAi*^ genotype where remarkable similarities with *pnr-Gal4* driven *UAS-Rab11*^*RNAi*^ were observed. *UAS-lgl*^*RNAi*^ alleles introgressed with *UAS-Rab11*^*RNAi*^ or *UAS-YFP-Rab11*^*Q70L*^ alleles were driven by *pnr-Gal4* and the number of flies which eclosed from the pupae were calculated for these introgressed alleles. The results were similar to the embryonic lethality experiments, where a rescue was observed in *pnr-Gal4* driven *UAS-Rab11*^*RNAi*^;*UAS-lgl*^*RNAi*^ individuals where out of an expected 100% driven flies, 29.48% eclosed, as compared to the *pnr–Gal4* driven*UAS-Rab11*^*RNAi*^ individuals where out of an expected 100%, 10.38% eclosed or *pnr-Gal4* driven *UAS-lgl*^*RNAi*^ individuals where out of an expected 100%, 12.22% eclosed. However, when *UAS-Rab11*^*Q70L*^; *UAS-lgl*^*RNAi*^ was driven with *pnr-Gal4*, the eclosion percentage of the driven individuals sharply fell to 4.54% out of an expected 100% driven progeny showing *Rab11* overexpression in an *UAS-lgl*^*RNAi*^ background results into augmentation of the lethal phenotype as compared to *pnr-Gal4* driven *UAS-lgl*^*RNAi*^ condition, corroborating our embryonic lethality assay results (Fig. 5c).

### The JNK mediated Dpp pathway is regulated by *Rab11* and *lgl* interaction during embryonic dorsal closure process

The JNK mediated Dpp signaling in the dorsolateral epithelium of stage 13-14 embryos of *Drosophila* is critical for the successful execution of the dorsal closure process. Inferring from our previous observations it can be safely stated that the genetic interplay of *Rab11* and *lgl* could be an important regulatory mechanism in this process which could further affect the JNK-Dpp pathway in the dorsolateral epithelial cells. In order to assess the effects of the *lgl*-*Rab11* interaction on the JNK signaling process, a transgenic JNK biosensor stock (*TRE-DsRed* also known as TRE-JNK) was duly introgressed with *pnr-Gal4*, thereby generating *TRE-JNK; pnr-Gal4* stock. As reported by Chatterjee and Bohmann, 2012, a transcriptional reporter expressing DsRED on activation of JNK pathway was constructed and inserted on the second chromosome by the ɸC31 based technique where the activation of c-JUN-FOS or AP1 transcription factor and its binding to TRE can be monitored by TRE (TPA responsive element) dependent DsRed expression, which provides an excellent system for monitoring JNK activity in the tissue under study and hence is an ideal bio-sensor of JNK signaling pathway.

Using this very biosensor, JNK activity was monitored in the background of different *UAS* constructs of *Rab11* and *lgl* driven by *pnr*^*MD237*^. It was observed that JNK patterns were indeed perturbed under different genetic backgrounds of *Rab11* and *lgl*. The improper expression patterns of JNK under the influence of different alleles could be a cause of the different morphological defects observed in each case. Not only JNK is altered differentially in different *pnr*-*Gal4* driven *UAS* constructs of *Rab11*^*RNAi*^ and *lgl*^*RNAi*^, the rescue which we observe in case of *pnr-Gal4* driven *UAS-Rab11*^*RNAi*^, *UAS-lgl*^*RNAi*^ individuals is also seen in terms of JNK signaling pattern, where both the *RNAi* driven individuals show an expression pattern similar to the wild type conditions whereas on the other hand, a *pnr-Gal4* driven *UAS-Rab11*^*CA*^; *UAS-lgl*^*RNAi*^ shows aggravated JNK signaling pattern (Fig. 6D-D’ & C-C’).

### *Rab11* and *lgl* mutants affect Dpp signaling (downstream of JNK) during dorsal closure in *Drosophila* embryos

JNK precedes Dpp signaling in the process of dorsal closure. Dpp or TGF-β is a morphogen essentially required for the regulation of downstream SMAD signaling which brings about cell morphology changes necessary to shape and guild tissue architecture at the time of development and differentiation. Thus in order to observe the effects of *Rab11* and *lgl* mutants in the JNK mediated Dpp signaling, we used a *Dpp-LacZ* reporter assay in order to assess the effects of these mutants in the process of dorsal closure. *Dpp-LacZ/CyO-Act-GFP; pnr*^*MD237*^/*TM3-Act-GFP* stock was used as a driver of *UAS-Rab11*^*RNAi*^; *UAS-lgl*^*RNAi*^ (A-A”), *UAS-lgl*^*RNAi*^ (B-B”) and *UAS-Rab11*^*RNAi*^ (C-C”) stocks, with undriven stock as control (D-D”). It was observed that, in controls (D-D”) a strong Dpp signaling occurs in the region of the leading edge cells which is sustained even after the completion of dorsal closure process as evident from the two parallel blue lines seen on the dorsal region of the embryo (D). However, this gets disrupted in *pnr-Gal4* driven *UAS-lgl*^*RNAi*^ or *UAS-Rab11*^*RNAi*^ conditions as observed in figures B and C, respectively, despite of the fact, a mild expression of Dpp exists at the time of dorsal closure process (B’-B” and C’-C”), though, not as robust as compared to the undriven controls (D’-D”). A somewhat rescue of the same is observed in *pnr-Gal4* driven *UAS-Rab11*^*RNAi*^; *UAS-lgl*^*RNAi*^ individuals (A-A”) which could be due to the restoration of the normal JNK signaling pattern as shown in the previous results.

## Discussion

The dorsal closure event signifies the gastrulation event in the fly embryos and is unique in the sense that it involves an extensive amount of coordinated cell shape changes exhibited by the contralaterally extending LE which finally zipper the dorsal opening of the germ band retracted embryo. As the LE extends, the constituent dorsal most and the dorsolateral cells undergo dorsoventral elongations which helps the lateral epidermis on either side extend and cover up the amnioserosa. Thus it is evident that the cell shape changes observed at the time of epithelial tissue extensions are highly polarized or directional. This property of the constituent cells is achieved on account of spatial cues which can be external, like the secreted morphogens; or internal, such as cytoplasmic morphogens or cell-polarity determining proteins. The cell-polarity determining proteins draw specific interest as they have been rigorously reported to be regulating intracellular cytoskeletal rearrangements and dynamics which is responsible for imparting cells their cognate shapes whereas on the other hand they also regulate critically important cell signaling pathways such as the JNK-Dpp pathway which in turn regulate the cytoskeletal dynamics as well as cell polarity. However, the mechanisms which intervene between cell polarity and cell signaling still remains an enigma even though an enormous amount of data has been generated in an attempt to join the two.

In the present study, we have tried to dissect a probable mechanism employed by the tumour suppressor, *lgl*, in developmental morphogenesis as has been reported by Manfruelli et al, 1996 where temperature sensitive, loss of function mutations of the gene show a characteristic dorsal open phenotype. This proves the instrumental role of the *lgl* in the process of wound healing which is popular as a component of the baso-lateral cell polarity complex along with Scribble and Dlg. However the cellular mechanisms which *lgl* regulates in order to accomplish the dorsal closure process yet remains to be answered although there are reliable evidences suggesting their profound effects on intra-cellular vesicle transport, cytoskeletal dynamics and cell signaling. In our experiments we find that *lgl* regulates spatio temporal expression pattern of Rab11, a small Ras like GTPase, which is essential for recycling cargo from Recycling Endosomes and the Trans Golgi Network to the plasma membrane. Reports of Sasikumar and Roy, 2009 and Mateus et al, 2011 suggest and demonstrate Rab11’s instrumental role in the dorsal closure process. The former group, i.e. Sasikumar and Roy have shown the effects of the loss of function allele, *Rab11*^*EP3017*^, in the DLE cells which fail to show proper cell morphologies along with the LE cells whereas Mateus et al, 2011 report its interaction with the early endosome Rab, Rab5 in the process of dorsal closure, where it is responsible for DE-Cadherin recycling in the LE cells at the time of dorsal closure.

These findings intrigued us to assess the interaction between *lgl* and *Rab11*, as their mutants show overlapping dorsal open phenotypes much like the mutants of the components of JNK pathway, like basket, slipper or puckered (Fig. 1). Reports of Zhang et al, 2005 and Langevin et al, 2005, suggest that *lgl* could be regulating polarized exocytosis by interacting with the components of the exocyst pathway, many of which also happen to be physical interactors of Rab11. Deriving cues from this information and exploiting the robust genetics of the *Gal4-UAS* system of targeted gene expression (Brand and Perrimon, 1993), a study on the effects of an ectopic perturbation of *lgl* in the DLE using *pnr-Gal4* (Kushnir et al, 2017) was made, where a down-regulation of *lgl* transcripts in *pnr-Gal4* driven *UAS-lgl*^*RNAi*^ individuals showed characteristic dorsal opening and puckering of embryonic cuticle as shown by *lgl*^*TS3*^ homozygotes reported by Manfruelli et al, 1996 (Fig. 1). Such cuticular aberrations appear due to inability of the cells of the LE to elongate and expand dorso-ventrally. This was further confirmed as we tracked these morphological changes in different genetic backgrounds of *pnr-Gal4* driven *UAS-lgl*^*RNAi*^ and *pnr-Gal4* driven *UAS-bsk*^*DN*^ where cells of the dorsolateral epithelium do not show the usual elongations as observed in an undriven background (Fig. 2). These cells are the ones which typically undergo JNK signaling responsible for the production and exocytosis of the Dpp morphogen (Zeitlinger et al, 1997 and Kushnir et al 2017) where Dpp in turn induces cell shape changes through its effects on the intracellular cytoskeletal dynamics. An interesting observation was made by Arquier et al, 2001, where *lgl* has been shown to regulate the emission of Dpp signal and thereby influencing dorsal closure process. Our observations in Fig. 7B-B” supports this finding but the mechanism via which *lgl* is able to regulate the release of Dpp morphogen yet needs to be addressed, although we prove here that a simultaneous knockdown of Rab11 in a *pnr-Gal4* driven *UAS-lgl*^*RNAi*^ background restores the Dpp expression pattern which is otherwise lost when *UAS-Rab11*^*RNAi*^ or *UAS-lgl*^*RNAi*^ are indivually driven by *pnr-Gal4*.

**Fig 1:**
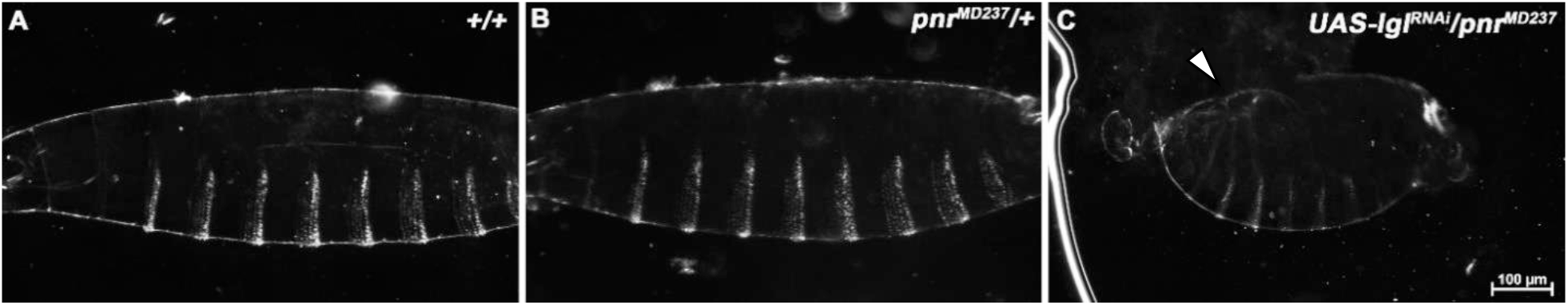
Dark field photomicrographs of embryonic cuticles under different genetic backgrounds, wild type (A), *pnr*^*MD237*^/+ heterozygous (B) and *pnr*^*MD237*^ driven *UAS-lgl*^*RNAi*^ (C). Note the shortening of the embryo, the puckering and the dorsal opening of the embryo (white arrow) of (C) as compared to (A) and (B).

**Fig 2.**
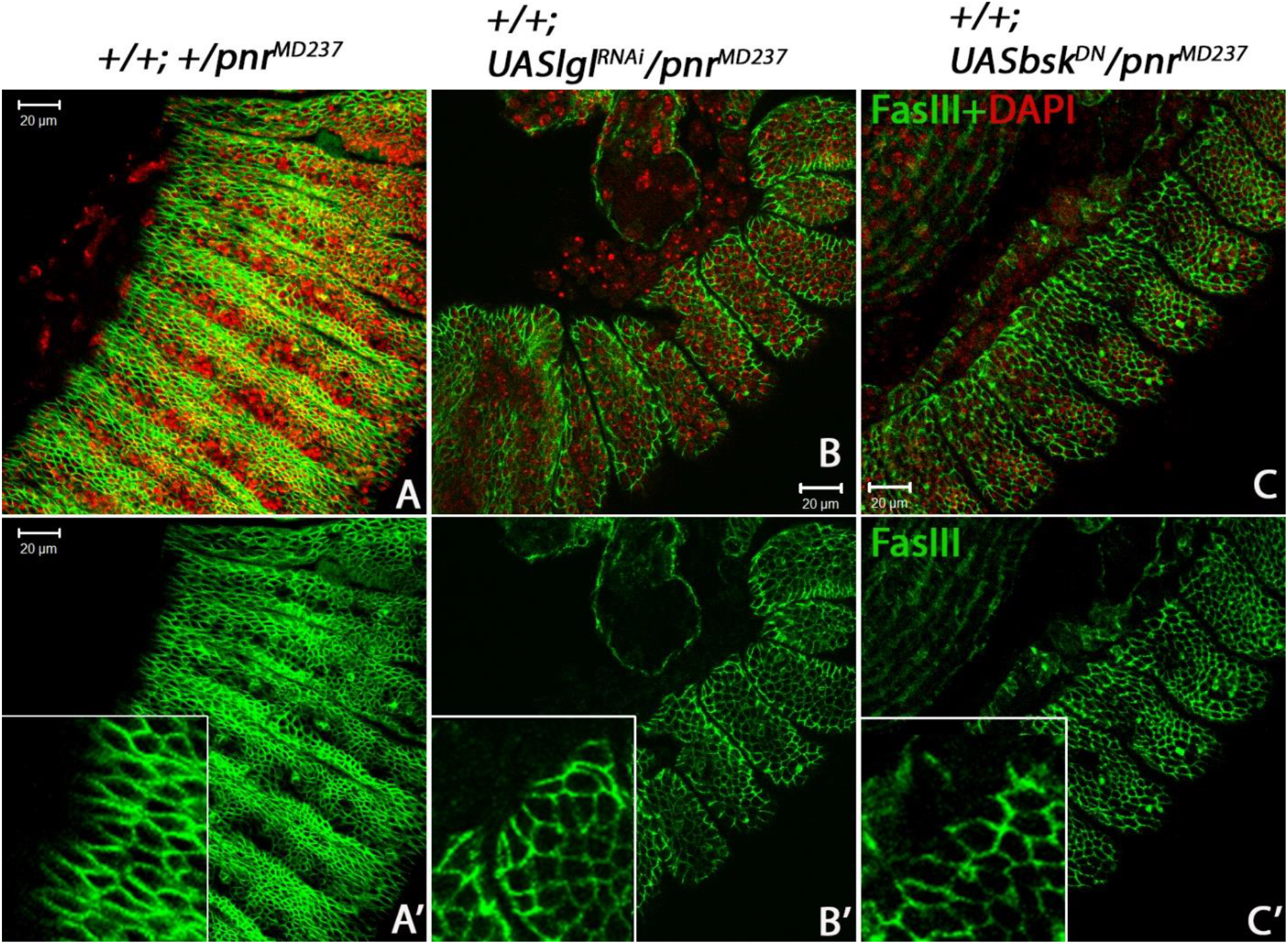
Confocal sections of late stage 13 embryos stained for FasIII (green) and DAPI (red) in different genetic backgrounds. Wild Type (A-A’) embryos are showing elongated epithelial cells which is a necessary morphological modification required to bring about dorsal closure. (B-B’ and C-C’) shows *pnr*^*MD237*^ driven *UAS-lgl*^*RNAi*^ and *UAS-bsk*^*DN*^ embryos, where cells of the dorso-lateral epithelium and the DLE cells do not show the regular dorso-ventrally elongated feature as shown by cells of the wild type embryos.

These observations as well as clues from available literature intrigued us to speculate into a probable role of *Rab11* in the functional aspects of *lgl* which it performs in the dorsal closure process where both *Rab11* loss of function mutants (Sasikumar and Roy, 2009) and *lgl* loss of function mutants show overlapping phenotypes. With standard genetic approaches, we indeed observed that *lgl* does synergize with *Rab11* in the dorsal closure process which we could trace at cellular levels (Fig 3 a). Our lethality assay results (Fig. 3b) also suggest that a strong interaction exists between *Rab11* and *lgl.* We observed that in a *pnr-Gal4* driven *UAS-lgl*^*RNAi*^ condition, 20.36% lethality was observed whereas on the other hand in a *pnr–Gal4* driven *UAS-Rab11*^*RNAi*^; *UAS-lgl*^*RNAi*^ background a lethality of 1.7% was observed which is a remarkable rescue. On the contrary, in a *pnr-Gal4* driven *UAS-YFP-Rab11*^*Q70L*^; *UAS-lgl*^*RNAi*^ background lethality values staggered up to 54.82% which suggests an aggravation of lethal phenotype. These observations could arise due to a regulatory effect of *lgl* on *Rab11* expression which has also been reported by Parsons et al, 2014 where *lgl*^−/−^ clones showed a characteristic accumulation of cytoplasmic Rab11. Our results in Fig. 4, corroborates this finding as in a *pnr-Gal4* driven *UAS-lgl*^*RNAi*^ background Rab11 expression levels get augmented in the dorso lateral epithelium with a consequent loss of the constituent cell shape and polarity. Interestingly, a Rab11 over-expression in a *pnr-Gal4* driven *UAS-YFP-Rab11*^*Q70L*^ embryos does not show a significant lethality which indicates the synergism and complementarities between the two loci required at the time of dorsal closure.

**Fig 3:**
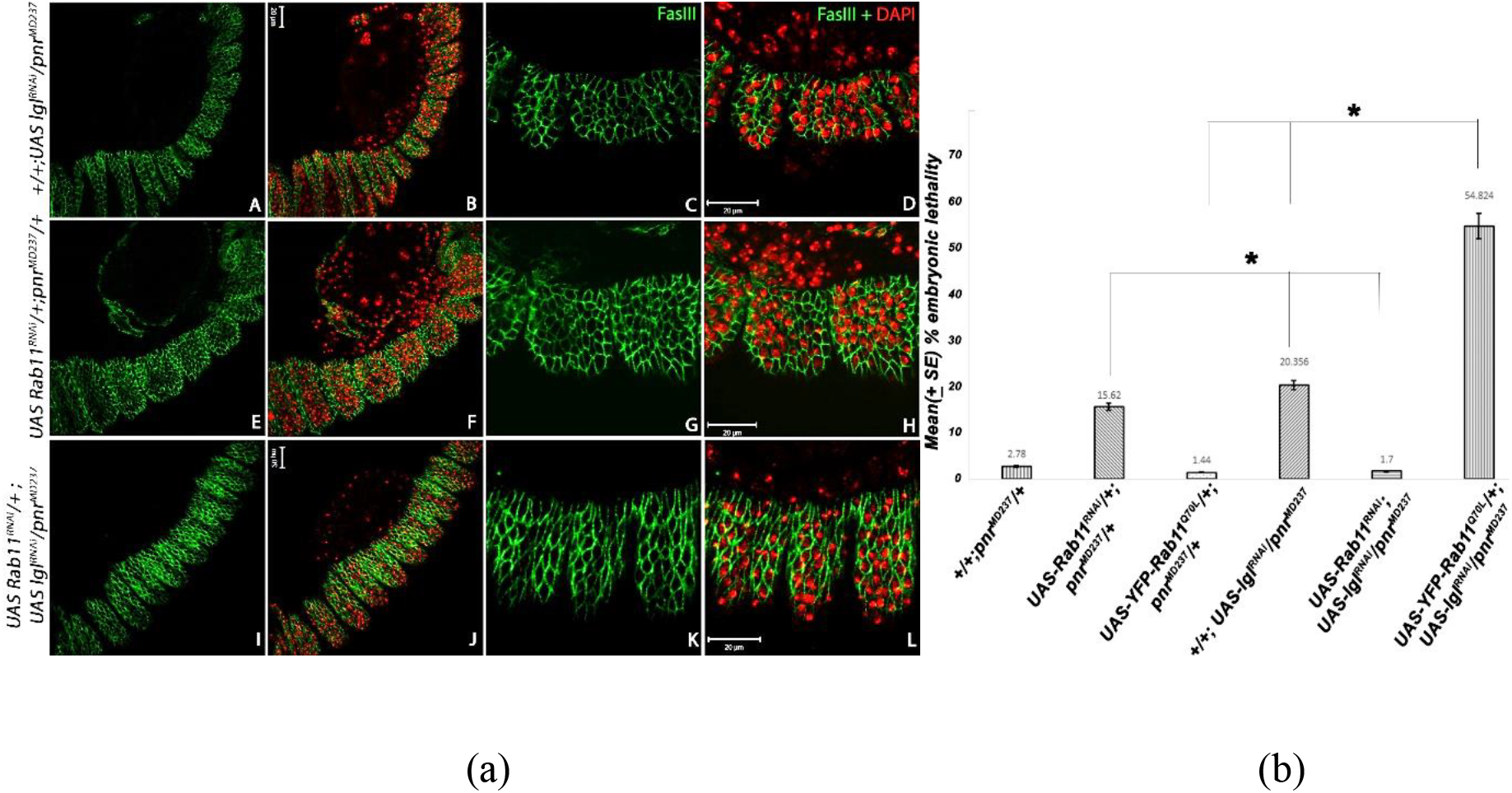
**(a)** Confocal sections showing lateral epithelial cells of stage 14 embryos immunostained for FasIII (green) and counterstained with DAPI (red pseudo-colour provided here). (A-D) represents *pnr-Gal4* driven *UAS-lgl*^*RNAi*^ individuals, where C-D are 2.5 times magnified sections of A and B, respectively. E-H represents *pnr-Gal4* driven *UAS-Rab11*^*RNAi*^ individuals where G and H are the 2.5 times magnified image of E and F, respectively. (I-L) represents *pnr-Gal4* driven *UAS-Rab11*^*RNAi*^; *UAS-lgl*^*RNAi*^ individuals, where K-L are 2.5 times magnified sections of I and J, respectively. The cell morphologies are much rescued in the panels I-L as compared to A-D and E-H. **(b)** Graph showing lethality caused by different *UAS* constructs of *Rab11* and *lgl* when driven by *pnr-Gal4*. The lethality values have been calculated out of an expected 100 percentage of *pnr* driven *UAS-Rab11* and *UAS-lgl* constructs. P value at ≤5% error has been considered significant.

**Fig 4.**
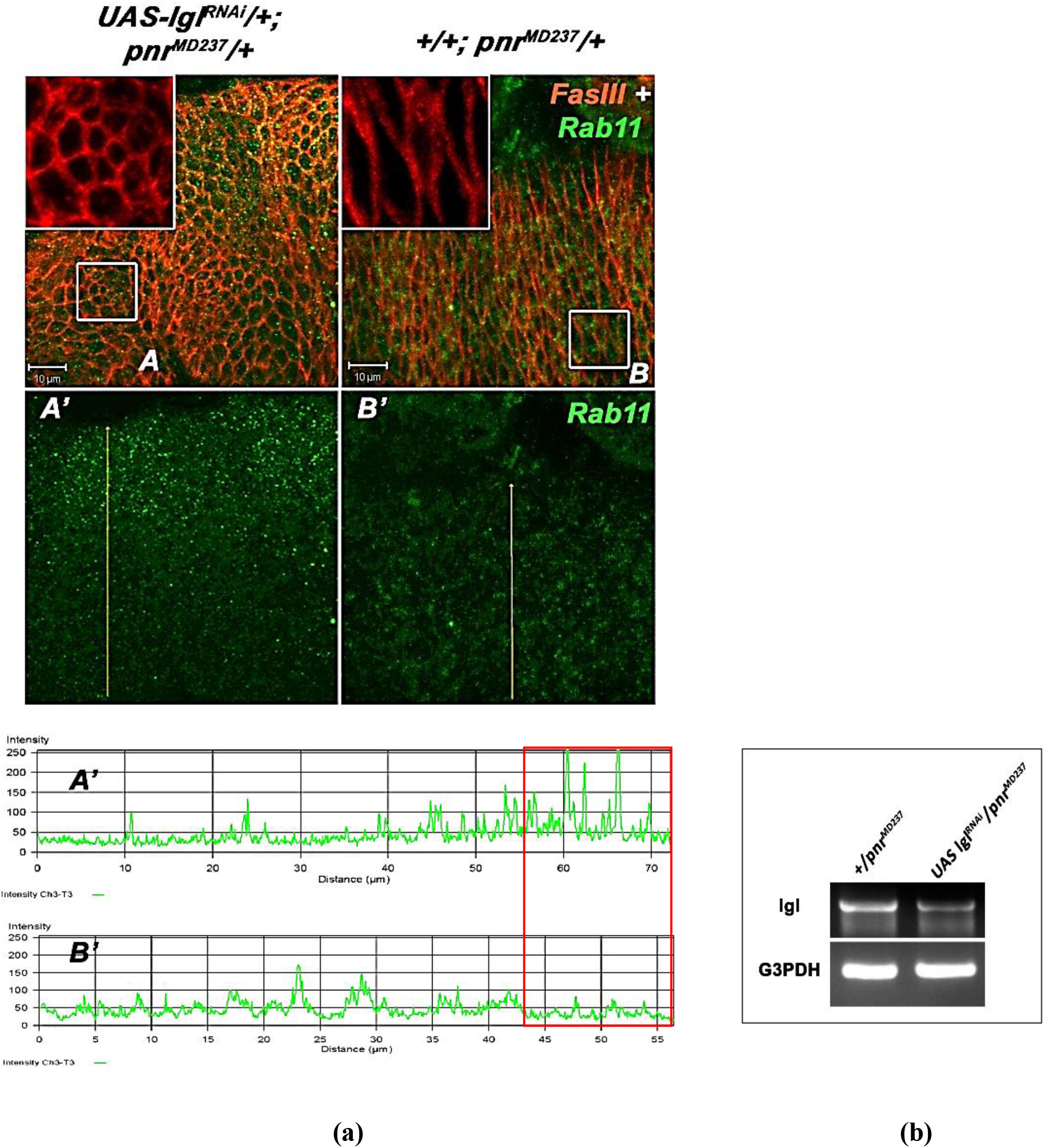
**(a)** Confocal projection images of late stage 13 *Drosophila* embryos immunostained for FasIII (red) and Rab11 (green). Insets in A and B represent the morphology of the lateral epithelial cells which undergo a strong loss of polarity as compared to the undriven control. Note the upsurge of Rab11 in the dorsolateral edge of the lateral epidermis which is consistent for all *pnr-Gal4* driven *UAS-lgl*^*RNAi*^ embryos. Quantitative graphical analysis of the mean intensities of Rab11 (green) through the regions of the yellow arrow reaching up to the dorsal end of the LE (A’ showing Rab11 expression levels in *pnr-Gal4* driven *UAS-lgl*^*RNAi*^ individuals and B’ showing Rab11 expression levels in an undriven condition). Note the upsurge of *Rab11* in A’ as compared to B’. **(b)** Semi-Quantitative RT –PCR results depicting the effect of *UAS-lgl*^*RNAi*^ driven by *pnr-Gal4* on *lgl* transcripts in stage 13 embryos where there is significant decline in *lgl* transcript levels.

An interesting observation was also made at the larval stages of *lgl* and *Rab11*introgressed flies (Fig. 5a), where individually driven (by *pnr-Gal4* or *pnr*^*MD237*^) *UAS-lgl*^*RNAi*^ or *UAS-Rab11*^*RNAi*^ larvae (the ones which do not undergo embryonic death) do not show dorsal lesions but perish as early embryos or pupae during thorax closure. However, the *pnr-Gal4* driven *UAS-Rab11*^*RNAi*^; *UAS-lgl*^*RNAi*^ individuals emerge in large numbers (Fig.5b) (justifying our embryonic lethality rescue experiments) amongst which a large fraction of larvae show characteristic dorsal lesions (Fig. 5 a) throughout the dorsal axis of the larvae. These lesions develop due to the partial rescue of dorsal closure defects which *UAS-Rab11*^*RNA i*^ or *UAS-lgl*^*RNAi*^ individually show when driven by *pnr-Gal4* (Fig. 5d). However, these larvae fail to develop into fully formed flies and show drastic thorax closure defects. The larvae which do not show these lesions develop into fully grown adults with properly formed thoraces which is also indicated from fly eclosion assay (Fig. 5c). A reason for this could be the different physiologies of the embryonic epithelium and the wing disc epithelium although they are executing parallel events of dorsal closure and thorax closure, respectively, governed by the same JNK-Dpp pathway. These observations suggest that *Rab11* and the tumour suppressor *lgl* interact in order to establish dorsal closure in the developing fly embryos. However the similarity of the genetic interaction between *lgl* and *Rab11* in both the dorsal closure and thorax closure processes is remarkable as both these events require a robust JNK-Dpp signaling at the migrating fronts of the contralaterally expanding epithelia.

**Fig 5.**
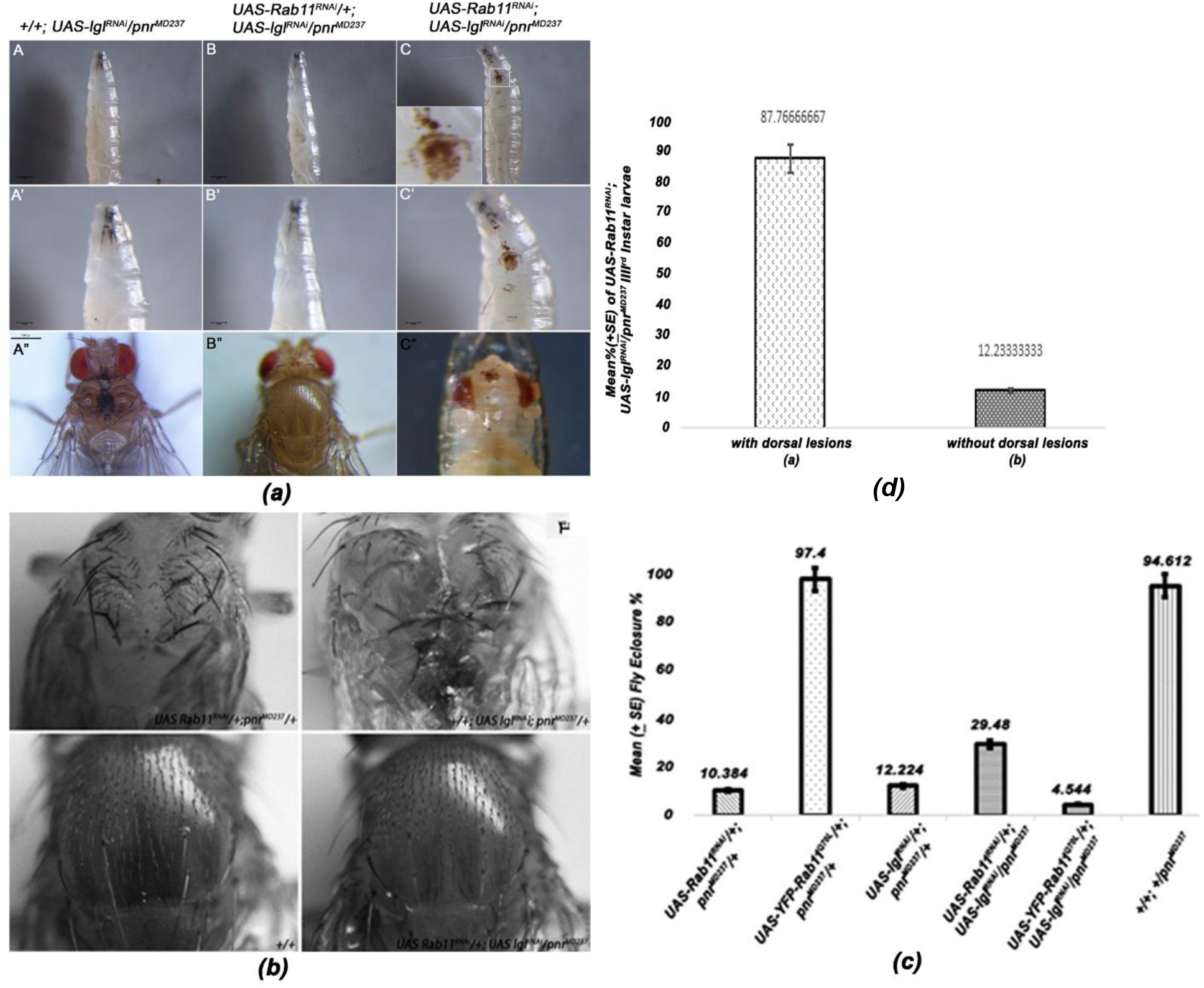
**a.** Photomicrographs of late third instar larvae showing dorsal cuticles of *pnr-Gal4* driven *UAS-lglRNAi*: (A-A”) completely rescued, *pnr-Gal4* driven *UAS-Rab11*^*RNAi*^;*UAS-lgl*^*RNAi*^ (B-B”) and partially rescued *pnr-Gal4* driven *UAS-Rab11*^*RNAi*^;*UAS-lgl*^*RNAi*^ individuals (C-C”). **b.** Photomicrographs of adult thoraces in a +/+ or wild type, *pnr-Gal4* driven *UAS-Rab11*^*RNAi*^, *pnr-Gal4* driven *UAS-lgl*^*RNAi*^ and *pnr-Gal4* driven *UAS-Rab11*^*RNAi*^;*UAS-lgl*^*RNAi*^ genetic backgrounds. **c.** Graphical representation of changes in mean percentage of fly eclosion as observed in *pnr-Gal4* driven *UAS-lgl*^*RNAi*^ background and *UAS-lgl*^*RNAi*^ introgressed with different *UAS*-constructs of *Rab11*. As observed earlier, *pnr-Gal4* driven *UAS-Rab11*^*RNAi*^, *UAS-Rab11CA* or *UAS-lgl*^*RNAi*^ show individual eclosion percentages of 10.38%, 97.4% and 12.22%. However, a mean eclosion percentage of 29.48% was observed in *pnr-Gal4* driven *UAS-Rab11*^*RNAi*^; *UAS-lgl*^*RNAi*^ progeny and on the contrary *pnr-Gal4* driven *UAS-YFP-Rab11*^*Q70L*^; *UAS-lgl*^*RNAi*^ shows an eclosion percentage of 4.54%. These flies showed severe thorax closure defects which suggests that *Rab11* overexpression in an *lgl* down-regulated genetic background results in an aggravated lethality due to dorsal closure and thorax closure defects. **d.** Graph representing the ratios of *pnr-Gal4* driven *UAS-Rab11*^*RNAi*^; *UAS-lgl*^*RNAi*^ larvae with and without dorsal lesions. Note that the ones with lesions are significantly higher than the ones without lesions.

As both the events of dorsal closure and thorax closure in *Drosophila* are carried out under the robust influence of JNK-Dpp expression (Agnes et al, 1999), it became imperative to analyze the effects of the genetic interaction of *Rab11* and *lgl* on the same. Therefore, in vivo reporters were resorted to in order to address this issue, where, TRE activated Ds-RED and LacZ reporter assays were employed to monitor the JNK and Dpp expression pattern in various genetic backgrounds of *Rab11* and *lgl* (Fig. 6 and Fig. 7). Reports of Escovar-Riesgo, 1997 and Fernandez, 2007, suggests that JNK–Dpp signaling promotes necessary coordinated cell shape changes required for dorsal closure which we could confirm in our observations (Supplementary figure 4 and 5). Significant alterations from the regular signaling pattern could be visible in the *pnr-Gal4* driven *UAS-lgl*^*RNAi*^ conditions (Fig 6-B’ and Fig. 7-B, B’ and B”) where JNK signaling as well as corresponding Dpp signaling is lowered as compared to the wild type conditions. JNK signaling shows a drastic down-regulation whereas Dpp signaling appears sparse as compared to the wild type. However, the rescue observed is significant as normal JNK and Dpp expression pattern is restored in a *pnr-Gal4* driven *UAS-Rab11*^*RNAi*^; *UAS-lgl*^*RNAi*^ genetic background (Fig. 6 D’ and Fig. 7 A). According to Arquier, 2001, *lgl* regulates the emission of Dpp morphogen in the embryonic epidermis. It could be that elevated Rab11 levels in *pnr-Gal4* driven *UAS-lgl*^*RNAi*^ background could interfere with the exocytosis of Dpp morphogen needed for necessary cell shape changes which when brought down to normal levels result in a regular pattern of Dpp signaling.

**Fig 6:**
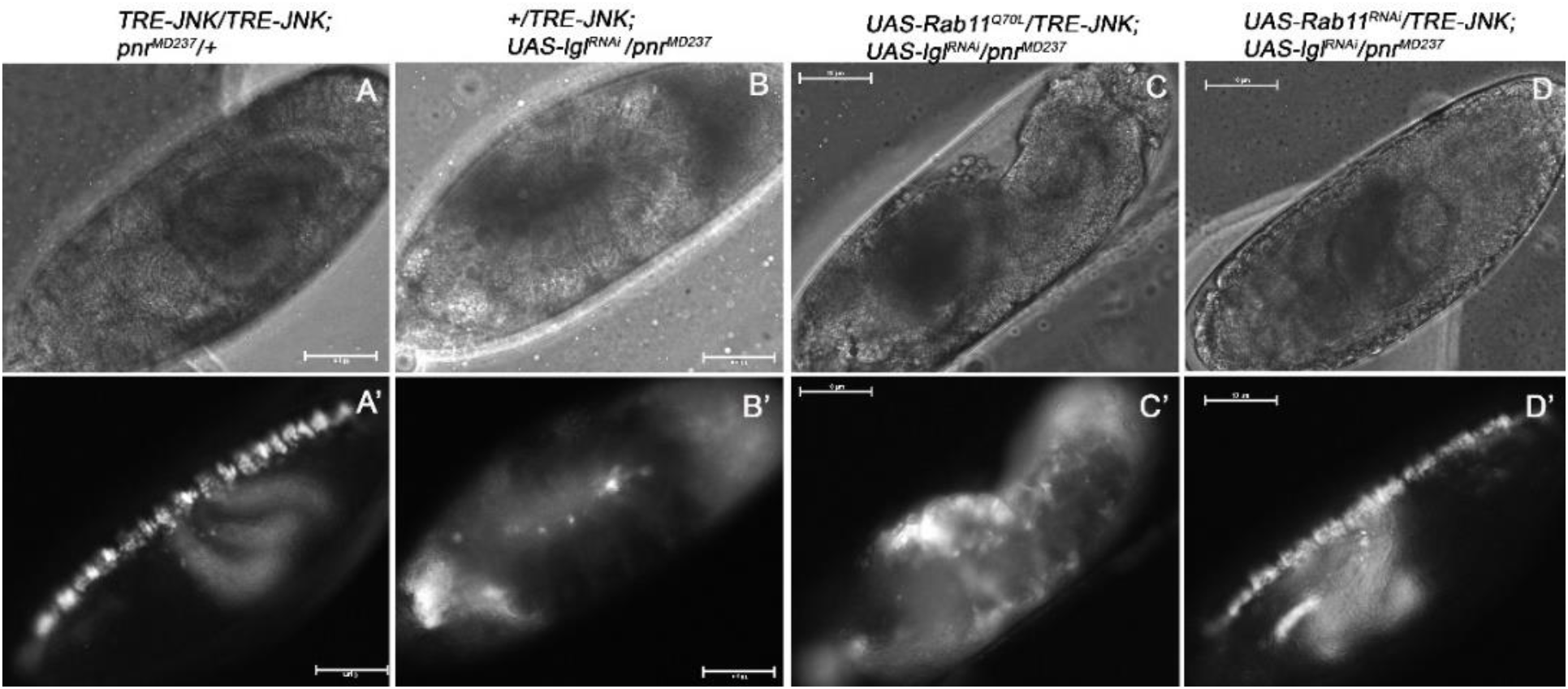
Phase contrast and Fluorescence images of 13-14h old *Drosophila* embryos in different genetic backgrounds of *pnr-Gal4*undriven (A-A’), *pnr-Gal4* driven *UAS-lgl*^*RNAi*^ (B-B’), *pnr-Gal4* driven *UAS-YFP-Rab11*^*Q70L*^; *UAS-lgl*^*RNAi*^ (C-C’) and *pnr-Gal4* driven *UAS-Rab11*^*RNAi*^;*UAS-lgl*^*RNAi*^ (D-D’)genetic backgrounds. 13-14 h developed embryos were observed which signifies the completion of the dorsal closure stage in all the above shown genetic backgrounds. Note the significant down-regulation of JNK signal in B’ as compared to A’ and D’. At the same time Rab11 when overexpressed in a *pnr-Gal4* driven *UAS-lgl*^*RNAi*^ background (C’), results in an upsurge of JNK signal signifying the effect of *Rab11* and *lgl* interaction on the JNK signaling pathway in the dorso-lateral epidermis of the developing embryos. The affected embryos in B’ and C’ show dorsal closure defects as reported in the earlier results.

**Fig 7:**
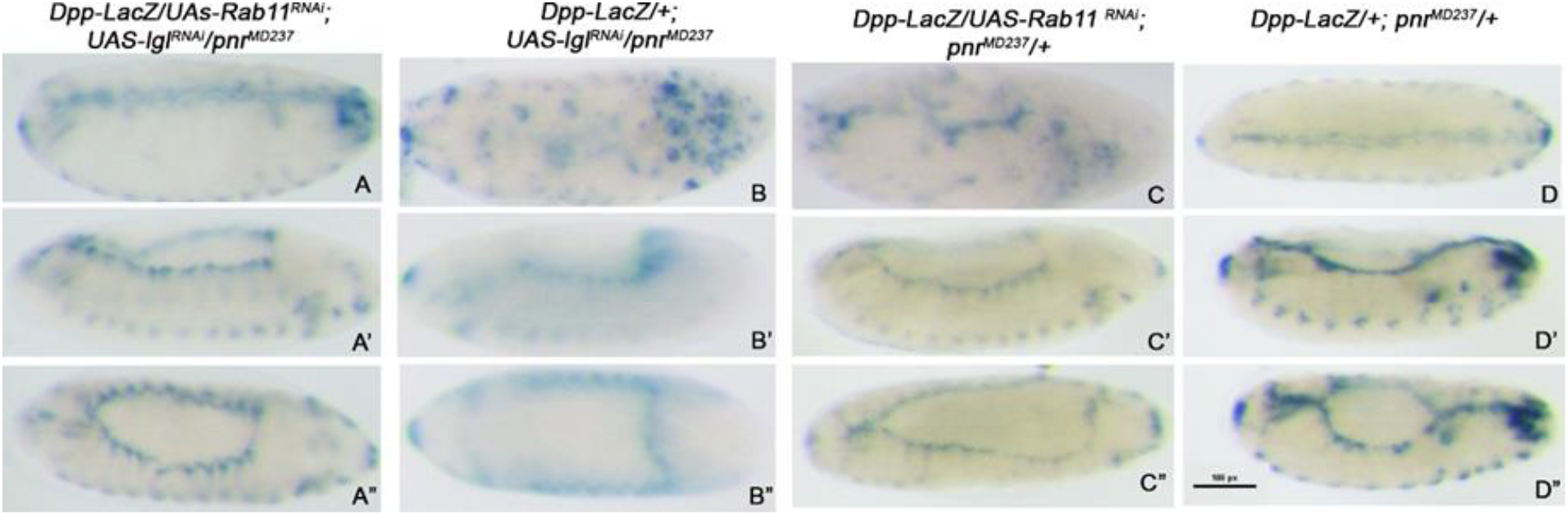
Bright field images of Lac-Z stained embryos under different genetic backgrounds, viz.,*pnr-Gal4* driven *UAS-Rab11*^*RNAi*^; *UAS-lgl*^*RNAi*^ (A-A”), *pnr-Gal4* driven *UAS-lgl*^*RNAi*^ (B-B”), *pnr-Gal4* driven *UAS-Rab11*^*RNAi*^ (C-C”) and undriven (D-D”) controls. The expression pattern of Lac-Z indicates Dpp expression pattern in the leading edge of the lateral epithelia of individual embryos. Observe the rescue in A-A” as compared to B-B” or C-C” panels which represents the restoration of proper Dpp signaling pattern on the down-regulation of *Rab11* in the *lgl* knockdown condition.

The above observations make it evident that Rab11 could be playing essential roles in *lgl* mediated epithelial morphogenesis. *lgl* mutations have been rigorously reported to be causing neoplastic epithelial tumours in *Drosophila* (Grifoni et al, 2013) and its roles in human cancers have been well documented by Schimanski et al, 2005 and Tsuruga et al, 2007. In their landmark review Schafer and Werner, 2008 suggest several parallels between the wound healing process and cancers which include TGFβ receptor induced SMAD signaling, which is also observed in our experiments. Using the dorsal closure of *Drosophila* as a model of wound healing, we could demonstrate that coordinated cell shape changes and contralateral fusion of epithelia requires the synergism between loci, which individually regulate cell polarity and intracellular recycling as we show the dorsal cuticular lesions of third instar larvae in a *pnr-Gal4* driven *UAS-Rab11*^*RNAi*^; *UAS-lgl*^*RNAi*^ background for the very first time (Fig. 5). Based on these observations it would further be interesting to dissect the mechanisms via which the tumour suppressor *lgl* could be regulating Rab11 expression pattern and consequent cell signaling of epithelial tissues, in order to exert its effects in developmental or diseased contexts.

## Acknowledgements

We thank the fly community for generously providing fly stocks. Special acknowledgements to Professor B. J. Rao for providing the TRE-JNK/CyO fly stock. We duly acknowledge the National facility for Laser Scanning Confocal Microscopy, Department of Zoology, Banaras Hindu University. The financial support of DST-FIST, UGC-UPE and CAS Zoology are duly acknowledged. We sincerely acknowledge the pioneering work of Dr. Satish Sasikumar, 2009, which suggested the role of Rab11 in epithelial morphogenesis. We sincerely thank University Grants Commission, New Delhi and Indian Council of Medical Research, New Delhi for providing the fellowships to NN.

## Supplementary Data 1 (S 1)

### Embryo selection scheme for immunofluorescence and expression analysis experiments

In order to distinguish between the *Gal4* driven and undriven embryos, *pnr-Gal4* was balanced with *TM3, Act-GFP,Ser1* and homozygous *UAS-lgl*^*RNAi*^ or *UAS-Rab11*^*RNAi*^; *UAS-lgl*^*RNAi*^ individuals were introgressed with this *Gal4* stock. The embryos so obtained were mounted in Halo-Carbon oil and observed under the fluorescence microscope (Nikon eclipse E800). A clear rescue of *pnr-Gal4* driven *UAS-Rab11*^*RNAi*^; *UAS-lgl*^*RNAi*^individuals along with the lethality of *pnr-Gal4* driven *UAS-lgl*^*RNAi*^ was observed.

**Fig. S1:**
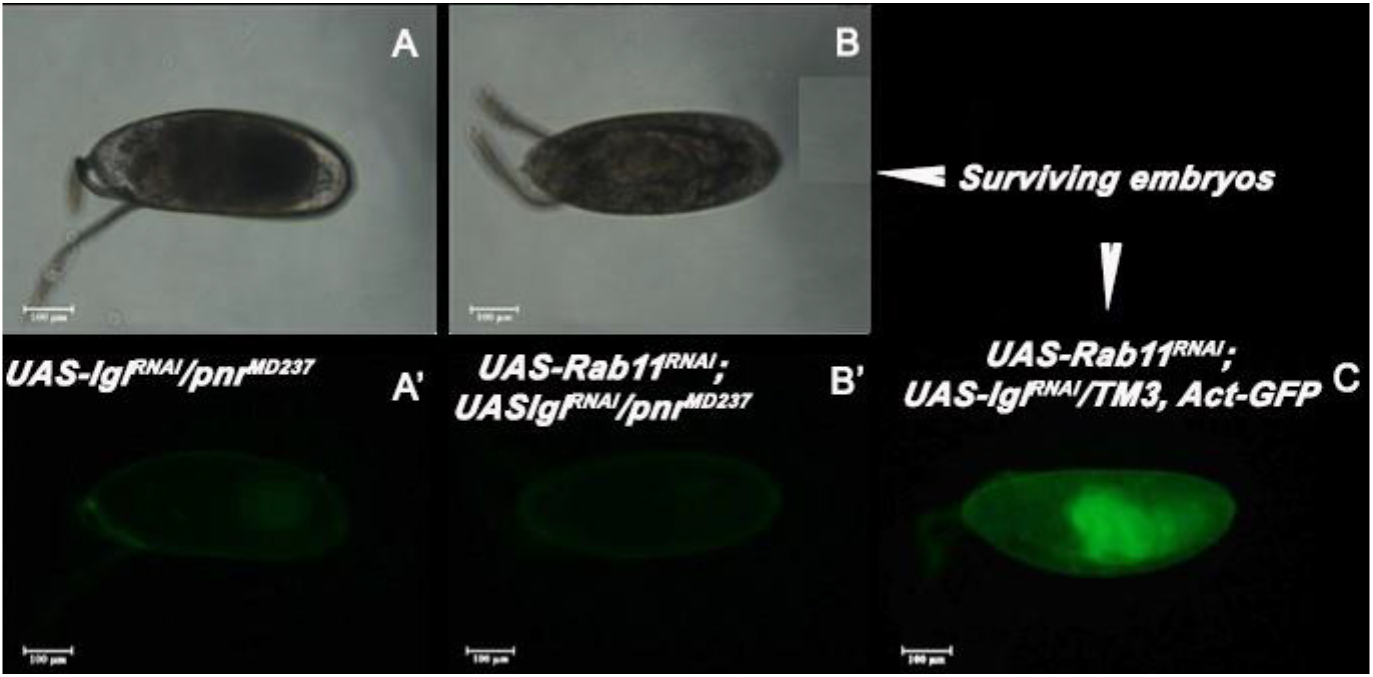
Phase contrast and Fluorescence images of 22-24 h developed fly embryos in *pnr-Gal4* driven *UAS-lgl*^*RNAi*^ background (A-A’), *UAS-Rab11*^*RNAi*^;*UAS-lgl*^*RNAi*^ background (B-B’) and balancer *TM3, Act-GFP, Ser1* containing embryos (C’). Note that the GFP negative embryos which are driven with *pnr-Gal4* show their respective phenotypes whereas the GFP positive embryos hatch out and continue on a normal developmental program.

## Supplementary Data2 (S2) For fig. 7

Lac Z reporter assay in undriven or wild type genetic background.

**Fig. S2:**
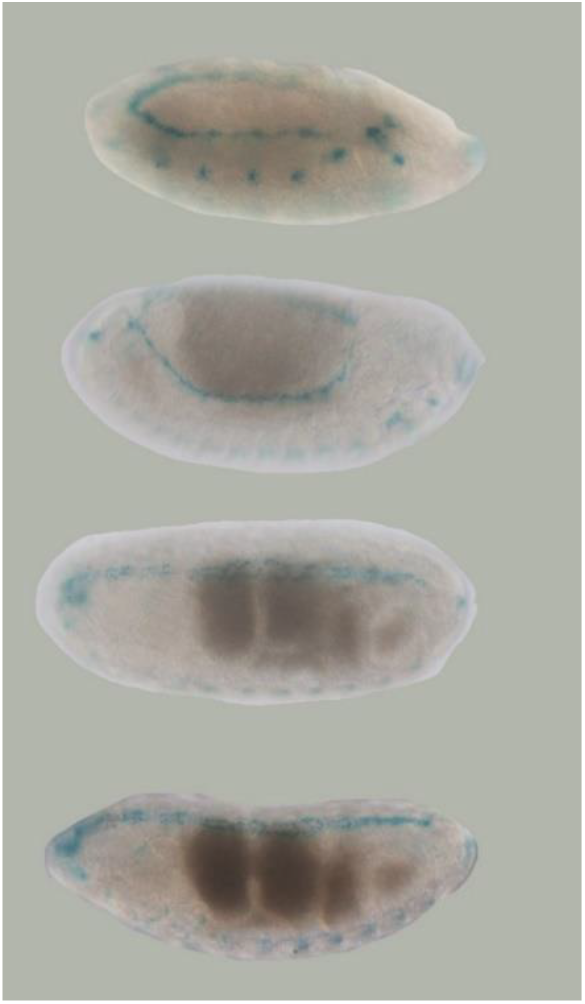
Lac Z reporter assay showing Dpp expression in different stages of development in a *pnr-Gal4/+* (undriven) genetic background. The same protocol has been used to stain the embryos shown in Fig. 7. Note the robust expression pattern of Dpp in the leading edge from the germ band extended to completion of dorsal closure stages.

## Supplementary data 3 (S3) For fig 6

JNK activity as seen through RFP expression in different stages of dorsal closure in a wild type genetic background.

**Fig. S3:**
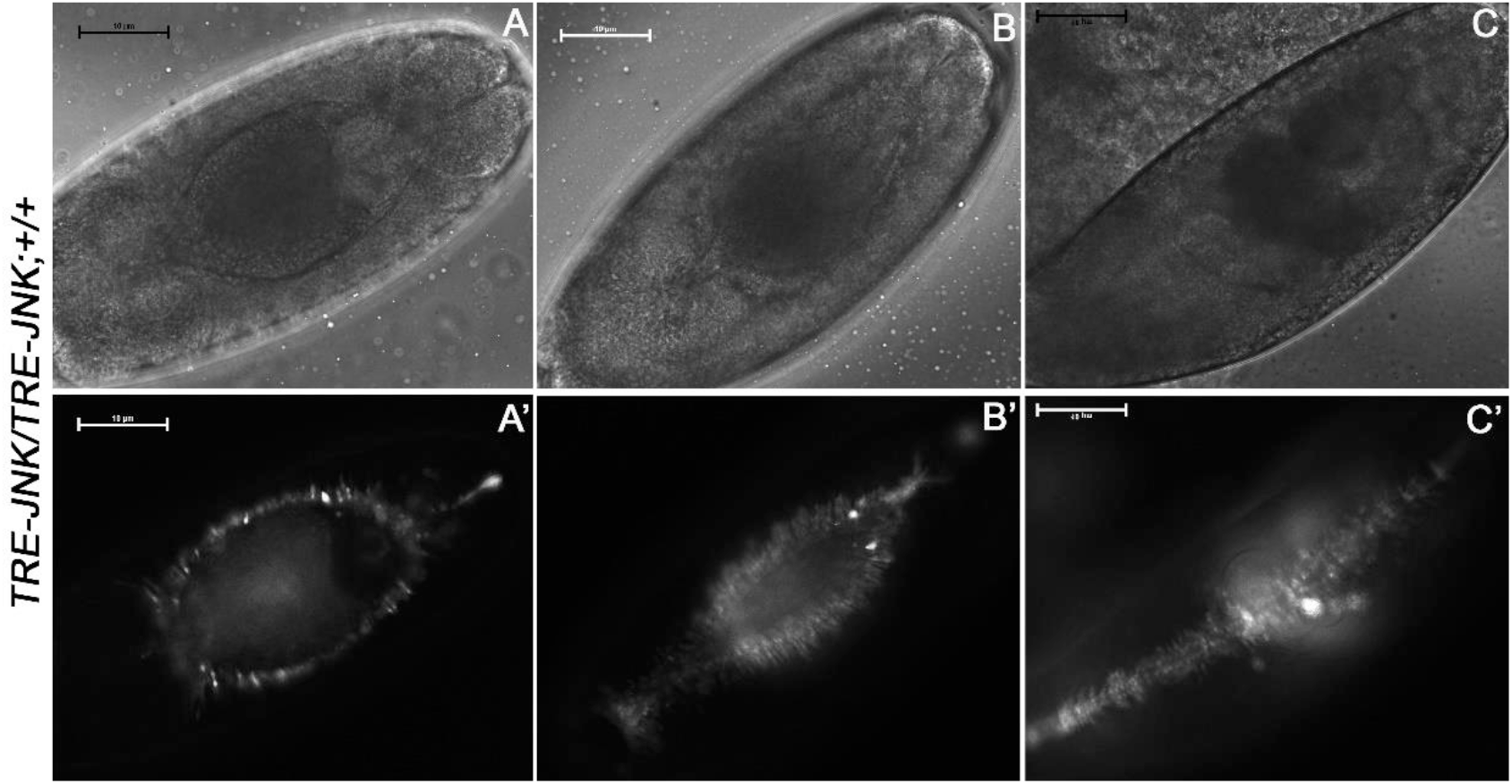
Phase contrast (A-C) and Fluorescence microscope (A’-C’) images of late 13, late14 and late 15 stages of TRE-JNK; + genotype embryos, showing JNK activity during the process of Dorsal closure. Observe the continual fluorescence expression in the cells of the Leading Edge and the dorsolateral epithelium which undergo morphological changes in order to complete the dorsal closure process.

## Supplementary data 4 (S4) demonstrating effects on Rab11 protein content in *pnr-Gal4* driven *UAS-Rab11*^*RNAi*^ individuals

**Fig. S 4:**
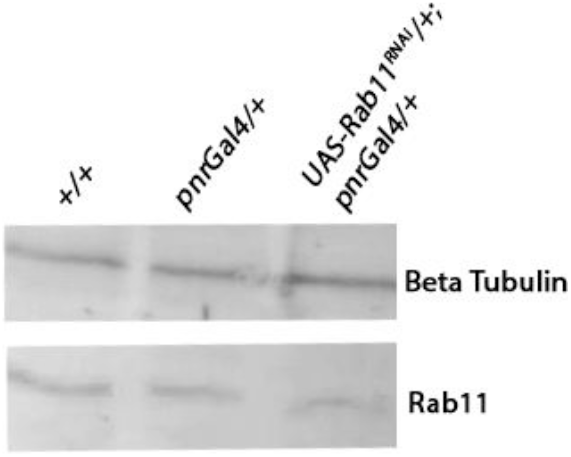
Western blot showing the effect of *pnr-Gal4* driven *UAS-Rab11*^*RNAi*^ on Rab11 protein levels. Note the decline in protein quantity in the third lane from left. This suggests the effective activity of the *UAS-Rab11*^*RNAi*^stock used in the experiment

**Supplementary table T1 in support of Fig 3b.**

| Cross scheme | Total fertilized | <u>Dead</u> | Total Death % (Dt) | Death% in controls (Due to balancers) (Dc) | Actual Death % in driven progeny (Da) | Fraction of driven progeny (According to Mendelian ratios) | Death % in 100% driven individuals |
| --- | --- | --- | --- | --- | --- | --- | --- |
| *pnr<sup>MD237</sup>/TM3,Ser*<br>X<br>+/CyO | 167 | 4 | 2.39(Control I) |  | 2.39 |  | 2.39 |
| *pnr<sup>MD237</sup>/TM3,Ser*<br>X<br>+/+ | 158 | 3 | 1.39(Control II) |  | 1.39 | 0.5 | 2.78 |
| *pnr<sup>MD237</sup>/TM3,Ser*<br>X<br>+;+/CyO-Tb | 109 | 62 | 56.39(Control III) |  | 56.39 | 0.25 | 56.39 |
| *pnr<sup>MD237</sup>/TM3,Ser*<br>X<br>*UAS-Rab11<sup>RNAi</sup>* | 402 | 37 | 9.2 | 1.39 (Control II) | 7.81 | 0.5 | 15.62 |
| *pnr<sup>MD237</sup>/TM3,Ser*<br>X<br>*UAS-Rab11<sup>Q70L</sup>/CyO* | 218 | 6 | 2.75 | 2.39 (Control I) | 0.36 | 0.25 | 1.44 |
| *pnr<sup>MD237</sup>/TM3,Ser*<br>X<br>*UAS-YFP-Rab11<sup>Q70L</sup>;UAS-lgl<sup>RNAi</sup>/CyO-Tb* | 418 | 293 | 70.09 | 56.39(Control III) | 13.71 | 0.25 | 54.82 |
| *pnr<sup>MD237</sup>/TM3,Ser*<br>X<br>*UAS-Rab11<sup>RNAi</sup>;UAS-lgl<sup>RNAi</sup>* | 491 | 11 | 2.24 | 1.39 (Control II) | 0.85 | 0.5 | 1.7 |
| *pnr<sup>MD237</sup>/TM3,Ser*<br>X<br>*UAS-lgl<sup>RNAi</sup>* | 389 | 45 | 11.57 | 1.39 (Control II) | 10.18 | 0.5 | 20.36 |

**Supplementary table T2 in support of fig. 5c.**

| <u>Cross scheme</u> | <u>Total</u><br><u>Flies</u><br><u>eclosed</u> | <u>Driven genotype</u> | <u>Total driven</u><br><u>progeny</u> | <u>Mean % of</u><br><u>driven progeny</u><br><u>which eclosed</u> | <u>Expected</u><br><u>% of</u><br><u>driven</u><br><u>progeny</u> | <u>Fraction of</u><br><u>driven progeny</u><br><u>(According to</u><br><u>Mendelian</u><br><u>ratios)</u> | <u>Fly</u><br><u>eclosure %</u><br><u>out of 10%</u><br><u>driven</u><br><u>progeny</u> |
| --- | --- | --- | --- | --- | --- | --- | --- |
| *pnr*<sup>MD237/TM3,Ser</sup><br>X<br>*Rab11RNAi* | 340 | *UAS-Rab11*<sup>RNAi/+</sup>;<br>*pnr*<sup>MD237/+</sup> | 16 | 5.19 | 50 | 0.5 | 10.38 |
| *pnr*<sup>MD237/TM3,Ser</sup><br>X<br>*UAS-YFP-*<br>*Rab11*<sup>Q70L/CyO</sup> | 605 | *UAS-YFP-*<br>*Rab11*<sup>CA/+</sup>;*pnr*<sup>MD237/+</sup> | 147 | 24.36 | 25 | 0.25 | 97.4 |
| *pnr*<sup>MD237/TM3,Ser</sup><br>X<br>*UAS-lgl*<sup>RNAi</sup> | 375 | *UAS-lgl*<sup>RNAi</sup>/*pnr*<sup>MD237</sup> | 22 | 6.11 | 50 | 0.5 | 12.22 |
| *pnr*<sup>MD237/TM3,Ser</sup><br>X<br>*UAS-*<br>*Rab11*<sup>RNAi</sup>;*UAS-*<br>*lgl*<sup>RNAi</sup> | 535 | *UAS-Rab11*<sup>RNAi</sup>;*UAS-*<br>*lgl*<sup>RNAi</sup>/*pnr*<sup>MD237</sup> | 79 | 14.74 | 50 | 0.5 | 29.48 |
| *pnr*<sup>MD237/TM3,Ser</sup><br>X<br>*UAS-YFP-*<br>*Rab11*<sup>Q70L</sup>;*UAS-*<br>*lgl*<sup>RNAi/CyO-Tb</sup> | 449 | *UAS-YFP-*<br>*Rab11*<sup>Q70L/+</sup>; *UAS-*<br>*lgl*<sup>RNAi</sup>/*pnr*<sup>MD237</sup> | 5 | 1.13 | 25 | 0.25 | 4.54 |
| *pnr*<sup>MD237/TM3,Ser</sup><br>X<br>+/+ | 316 | +/ *pnr*<sup>MD237</sup> | 310 | 47.30 | 50 | 0.5 | 94.61 |

